# *PHASIS*: A computational suite for *de novo* discovery and characterization of phased, siRNA-generating loci and their miRNA triggers

**DOI:** 10.1101/158832

**Authors:** Atul Kakrana, Pingchuan Li, Parth Patel, Reza Hammond, Deepti Anand, Sandra M. Mathioni, Blake C. Meyers

## Abstract

Phased, secondary siRNAs (phasiRNAs) are found widely in plants, from protein-coding transcripts and long, non-coding RNAs; animal piRNAs are also phased. Integrated methods characterizing *“PHAS”* loci are unavailable, and existing methods are quite limited and inefficient in handling large volumes of sequencing data. The *PHASIS* suite described here provides complete tools for the computational characterization of *PHAS* loci, with an emphasis on plants, in which these loci are numerous. Benchmarked comparisons demonstrate that *PHASIS* is sensitive, highly scalable and fast. Importantly, *PHASIS* eliminates the requirement of a sequenced genome and PARE/degradome data for discovery of phasiRNAs and their miRNA triggers.

## Background

Phased siRNAs (phasiRNAs) are a major subclass of secondary siRNAs, found extensively in plants [1]. The defining characteristic of phasiRNAs is the DCL-catalyzed processing of doublestranded RNA (dsRNA) precursors, starting from a precisely delimited 5’ terminus and generating regularly-spaced 21-or 24-nt populations of siRNAs [2]. PhasiRNAs can be further subdivided into three main categories based on their precursor mRNAs and spatiotemporal patterns of accumulation: i) The first phasiRNAs identified, so-called *trans*-acting siRNAs (tasiRNAs) generated from a small set of long, non-coding mRNAs (lncRNAs) referred to as *TAS* genes [3-5]; ii) phasiRNAs from protein-coding transcripts, such as *NB-LRRs* or *PPRs* [6]; and iii) two classes, 21-nt premeiotic or 24-nt meiotic phasiRNAs, highly enriched in reproductive tissues and also produced from lncRNAs, reported in grasses, but with no as-yet reported targets [2,7]. Thus, the umbrella name of “phasiRNAs” refers simply to their biogenesis and not their function (unlike the subset of tasiRNAs) because many phasiRNAs lack validated targets, either in *cis* or *trans* [6,8].

The biogenesis of phasiRNAs in plants is dependent on a triggering mechanism that sets the phase of the resulting secondary siRNAs, generated from a specific nucleotide in the mRNA precursor. To date, the only described type of trigger is a miRNA, and a breakthrough in our understanding of plant miRNA function came with the observation that all or nearly all 22-nt miRNAs trigger phasiRNA biogenesis from their targets [9,10]. These miRNA triggers function via the ARGONAUTE (AGO) proteins into which they are loaded, and since phasiRNA biogenesis requires both SGS3 and RDR6 [3,4], there may be interactions between these proteins, ultimately recruiting DCL4 or DCL5. SGS3 and RDR6 proteins function in the cytoplasm, forming siRNA bodies [11]. Recent work has identified membrane-bound polysomes in the rough ER as the site where miRNA triggers of phasiRNAs accumulate, leading to phasiRNA biogenesis [12]. miRNA triggers are thus an important component in the analysis of plant phasiRNAs, and the identification of specific triggers with specific *PHAS* targets is an integral part of phasiRNA analysis.

Since the discovery of phasiRNAs in 2005, *TAS* genes have been characterized in detail, especially the eight loci in Arabidopsis but these represent only a small fraction of the *PHAS* repertoire found in many plant genomes. In other eudicot genomes, there are hundreds of protein-coding genes that are targeted by diverse miRNAs, many of which are lineage-specific [8,13-15]. Grass genomes contain even more *PHAS* loci. For example, loci yielding reproductive phasiRNAs number in the hundreds to thousands in maize [7] and rice [16], and have yet to be characterized broadly in monocots or other lineages outside of the grasses. These include the premeiotic *PHAS* loci that are targeted by miR2118 family members, triggering production of 21-nt phasiRNAs accumulating in early anther development, and the 24*-PHAS* loci that are targeted by miR2275 family members, triggering production of 24-nt phasiRNAs, accumulating in anthers during meiosis [7]. Analysis of the spruce genome, a gymnosperm that speciated ~325 million years before the evolution of monocots and eudicots, identified over 2000 *PHAS* loci mostly from protein-coding genes, including over 750 *NB-LRRs* [17]. Thus, plant *PHAS* loci are widely prevalent and highly variable from genome to genome both in the total number and in terms of the types of loci that generate them. Characterization of *PHAS* loci from each sequenced plant genome will provide insights into this unusual type of post-transcriptional control, its evolution, and diversification.

Tools for the *de novo* identification of *PHAS* genes (or loci) to date have required an assembled genome for their discovery and additional experimental data such as PARE [18], degradome [19] or GMUCT [20] libraries to identify their miRNA triggers. Integrated tools for discovery and in-depth characterization of *PHAS* genes have not yet been developed, and the existing options are both limited in number and function. These algorithmic limitations and bioinformatic gaps along with the increasing depth and volume of sequencing data necessitates scalable, fast and advanced methods to study this relatively new class of secondary siRNAs for which parallels exist between plants and animals [2,13,21,22]. Motivated by this need for software, the prospect of discovering novel phasiRNA modules, the emerging importance of phasiRNAs, and the explosion in the number of plant species that are being investigated for small RNAs (sRNAs), we developed a new computational suite that we call *“PHASIS”*. The name *“PHASIS”* is from the ancient Greek city of Phasis, a destination for Jason and the Argonauts according to Greek mythology; we selected the name as it links the colloquialism “*phasis*” as short for phasiRNAs, with the Argonaut proteins that bind them. This set of tools facilitates the discovery, quantification, annotation, comparison of *PHAS* loci (and precursors) and identification of their miRNA triggers, from a few to hundreds of sRNA libraries in a single run. *PHASIS* not only addresses crucial bioinformatic gaps while providing an integrated and flexible workflow for the comprehensive study of *PHAS* loci, but it is also fast and sensitive.

## Results

### Assessment and benchmarking

We first sought to assess the sensitivity and specificity for *PHASIS;* ideally, this would be done with a gold-standard reference set of experimentally-validated *PHAS* loci in plants. While the definition of “gold standard” is as-yet unclear for *PHAS* loci, the recently-described maize loci are among the most exhaustively characterized [7], and thus we used these data below. We also compared *PHASIS* predictions and performance with PhaseTank [23]. Currently, two computational tools are capable of *de novo* discovery of *PHAS* loci – PhaseTank [23] and ShortStack [24]. *PhaseTank* is exclusively built for predicting *PHAS* loci in plants, while *ShortStack* aims to annotate and quantify diverse sRNA-associated genes (or clusters), and it's typically deployed for characterizing miRNAs in plants and animals [24]. A direct comparison between *PHASIS* and *ShortStack* is not possible due to significant differences in their scope, utility and workflow (Table 1). So, for comparative benchmarking, we chose *PhaseTank*, mainly because of matching objectives and its published superiority over *ShortStack* in predicting *PHAS* loci [23]. Benchmarking was performed across five plant species – *Arabidopsis thaliana* (Arabidopsis), *Brachypodium distachyon* (Brachypodium), *Oryza sativa* (rice), *Zea mays* (maize) and *Lilium maculatum* (Lilium). These species were selected based on availability of high-quality nuclear genome assemblies or anther transcriptomes (in case of Lilium – generated for a different study but included here), and deep sRNA libraries from premeiotic and meiotic anther or from at least one of these two stages that should contain many reproductive phasiRNAs **(Supplementary Table** 1). Arabidopsis was included because it was originally used in *PhaseTank* benchmarking [23]. For *PhaseTank*, the reference genome, transcriptome and sRNA libraries were converted to the appropriate formats, and the time for file conversion process, although complex and lengthy, was not added in the *PhaseTank* runtimes. *PHASIS* and *PhaseTank* use inherently different scoring schemas; because of this difference, we used a conservative p-value (1e-05) for *PHASIS* and the recommended score (i.e. 15) for *PhaseTank*. All benchmarks were performed on a 28 core, 2.42 GHz machine with 512 GB of RAM, running CentOS 6.6.

**Table 1.** Comparison of features from existing tools for phasiRNA characterization.

|  | Software & citation: | *ShortStack*,<br>Axtell MJ<br>(2013) | *PhaseTank*,<br>Guo et al.<br>(2015) | *PHASIS* |
| --- | --- | --- | --- | --- |
| *PHAS* prediction-related features | Tool-specific data format requirement? | no | yes | no |
|  | Library-specific results? | no | no | yes |
|  | *PHAS* prediction in w/o genome assembly? | no | no | yes |
|  | Results grouped based on stage, tissue or treatments? | no | no | yes |
|  | *PHAS* comparison between groups? | no | no | yes |
|  | *PHAS* locus annotation? | yes (from genome GFF only) | no | yes |
|  | *PHAS* trigger prediction w/o PARE data? | no | no | yes |
| Features unrelated to *PHAS* prediction | miRNA/hairpin locus prediction? | yes | no | no |
|  | Whole-genome report of sRNA cluster characteristics? | yes | no | no |

### PHAS prediction and runtime performance

We first compared *PHAS* loci and transcript predictions from *PHASIS* and *PhaseTank*. Since Arabidopsis lacks 24-*PHAS* loci (none have ever been published, nor have we found any), and the genome encodes just eight *TAS* genes, these were excluded from quantification of prediction and speed comparisons. *PHASIS* demonstrated an edge over *PhaseTank* in *PHAS* predictions: in genomic analyses, it predicted up to 2.5 times more *PHAS* loci, ranging from 73 24-*PHAS* (145% gain) to 380 21*-PHAS* (24% gain) loci in Brachypodium and rice respectively **(Table 2).** The biggest gain was observed in an analysis of the Lilium transcriptome, in which *PHASIS* predicted ~10 times (n=408) more 21*-PHAS* and 18 times (n=9065) more 24-*PHAS* precursor transcripts compared to *PhaseTank* (Figure 2). The specific data format requirements of *PhaseTank* made it difficult to accurately determine the set of common *PHAS* predictions (the ‘common *PHAS* pool’, hereafter) for transcriptome level analysis, however, by matching the sequences we determined that *PHASIS* captured at least 66% of 21*-PHAS* and 99% of 24-*PHAS* predictions from *PhaseTank*. For genomic analyses, *PHASIS* captured >80% of *PhaseTank* predictions, except in rice and Arabidopsis in which *PhaseTank* predicted additional 24-*PHAS* loci **(Table 2).**

**Table 2.** Comparison of predictions for *PHAS* loci, precursor transcripts, and their miRNA triggers.

| Species | Type | *PHAS* locus gain with *PHASIS* over *PhaseTank* | *PhaseTank* *PHAS* loci captured by *PHASIS* | Gain in miRNA triggers: *PHASIS* (PARE supported) vs. *PhaseTank* (PARE supported) | Gain in miRNA triggers, (predicted) *PhaseTank* support |
| --- | --- | --- | --- | --- | --- |
| Arabidopsis | 21-PHAS | 21% | 84% | -54% | -18% |
| Brachypodium | 21-PHAS | 145% | 79% | 76% | 178% |
|  | 24-PHAS | 49% | 85% | 36% | 69% |
| Rice | 21-PHAS | 24% | 97% | 5% | 54% |
|  | 24-PHAS | -33% | 29% | No predictions | N.D. |
| Maize | 21-PHAS | 81% | 97% | 4% | 55% |
|  | 24-PHAS | 59% | 86% | 9% | 64% |
| Lilium | 21-PHAS | 907% | 67%* | No PARE data | No PARE data |
|  | 24-PHAS | 1694% | 94%* | No PARE data | No PARE data |

For Lilium, no PARE data were available for trigger predictions. N.D. = not determined, capture trigger miRNAs.

The additional 24-*PHAS* loci predicted by *PhaseTank* in rice and Arabidopsis all had significantly lower quality scores (from *PhaseTank)* compared to the common *PHAS* pool, as did the *PhaseTank-exclusive 21-* and 24-*PHAS* predictions from other species. The average quality scores computed for each species were 1.7 to 7.8 times lower compared to the common *PHAS* pool (p-value < 0.001, t-test) **(Supplementary Table 2);** thus, the predictions exclusive to *PhaseTank* are likely unphased and a misinterpretation of loci yielding profuse heterochromatic siRNAs (hc-siRNAs). This may explain the 24-*PHAS* predictions in Arabidopsis by *PhaseTank* (Figure 2b and Table 2), as 24-nt phasiRNAs have not been reported in Arabidopsis despite exhaustive analyses [1]. Nonetheless, considering that these *PhaseTank* predictions could represent weak *PHAS* loci, we attempted to capture them by running *PHASIS* at lower p-value cutoff (1e-03) but still failed to detect >96% of them. Manual investigation of a portion of these *PHAS* loci using our custom sRNA browser, which uses a slightly different *PHAS* scoring schema [25], revealed that these are indeed either unphased or show typical characteristics of hc-siRNA loci such as similarity to transposons, and we concluded that these are false positives predicted by *PhaseTank* (Supplementary Figure 1A). However, we could detect 70% (n=67) of the total *24-PHAS PhaseTank* predictions in rice at the lower p-value cutoff (1e-03) of *PHASIS*, and a majority of these showed weak phasing patterns **(Supplementary Figure 1B),** suggesting that *PHASIS* missed these at the selected cutoff. However, the count of 24-*PHAS* loci predicted in rice by both tools in these libraries from a recent study [16], was lower than earlier estimates [2], indicating that the libraries likely missed meiotic peak of accumulation. These contrasting observations – Arabidopsis, in which *PHASIS* correctly excluded 24-*PHAS* predictions even at relaxed cutoff, versus rice, in which it correctly captured 70% of weakly phased 24-*PHAS* loci – highlights differences in scoring in the two tools, with the default *PHASIS* p-value cutoff (1e-05) more stringent than that of *PhaseTank* (score=15). Using a lower p-value cutoff for *PHASIS* could further increase the gain in *PHAS* predictions over *PhaseTank* without adding much noise.

**Figure 1.**
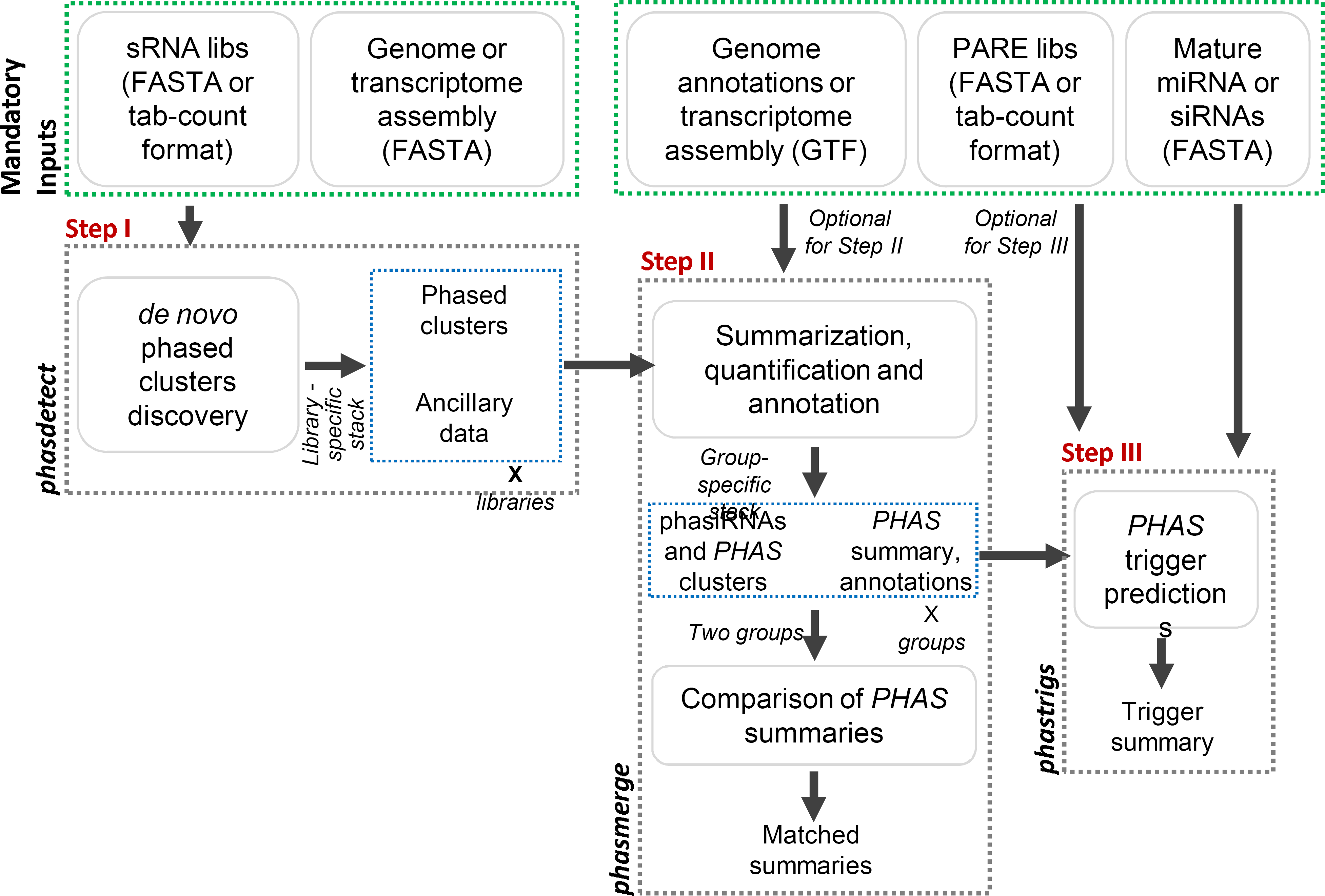
PHASIS workflow. *PHAS* loci or precursors transcripts are predicted through *phasdetect* in the first step. The library-specific list of *PHAS* predictions can be summarized and annotated through *phasmerge* for libraries of interest into a *PHAS* summary. These summaries from two different groups can also be compared using “compare” mode of *phasmerge*. Triggers for *PHAS* summaries are identified through *phastrigs* either with PARE data in “validation” mode or without any experimental data in “prediction” mode. Selection between these two modes is made automatically based on a PARE library input or the lack of it. All analysis steps are independent and their execution depends upon the requirements of the user.

**Figure 2.**
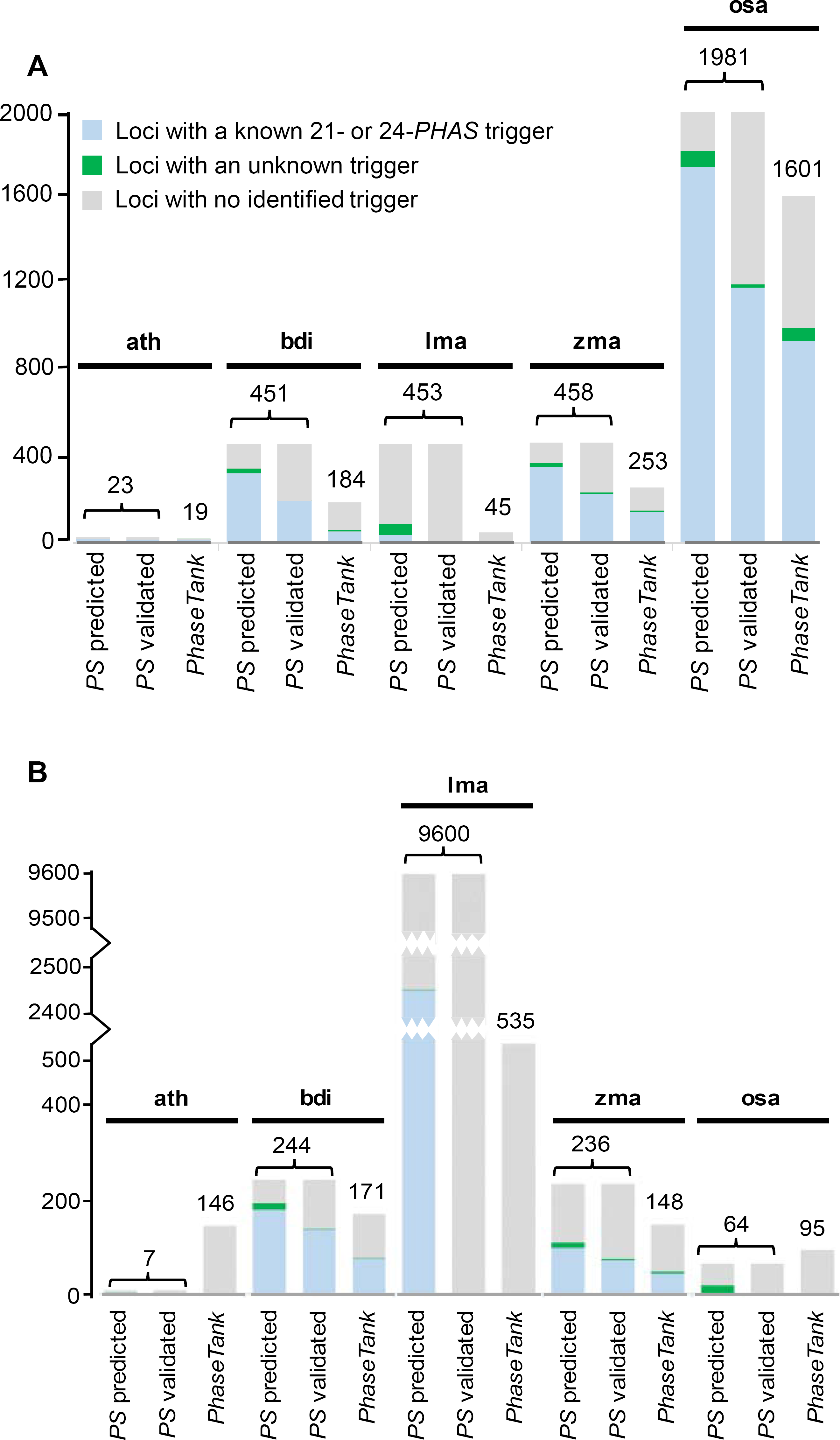
Number of *PHAS* loci or transcripts and their triggers, predicted by *PHASIS*. *PHASIS* is labelled as *'PS'* and it is compared to *PhaseTank* for benchmarking. **A**) 21*-PHAS* and **B**) 24-*PHAS* loci identified by both tools along with their triggers in Arabidopsis (ath), Brachypodium (bdi), Lilium (Ima), rice (osa) and maize (zma). For *PHASIS* trigger prediction, results from both “validation” and “prediction” mode were included. The bars for Lilium 24-*PHAS* loci are split at two different points for display purposes. Triggers assigned to PHAS loci that do not match with known or published miRNA triggers were represented as ‘unknown' triggers.

We manually investigated 21-and 24*-PHAS* predictions that are exclusive to *PHASIS*, using our public, custom genome browser (https://mpss.danforthcenter.org/). The majority of these displayed characteristics matching those of the canonical 21-and 24*-PHAS* loci reported in maize [7] **(Supplementary Figure 2).** Moreover, a major proportion of these *PHASIS*-exclusive predictions had PARE-validated miRNA triggers, matching to the earlier reports from maize, rice and Brachypodium [2,7,13]. Next, we compared prediction runtimes of *PHASIS* and *PhaseTank* from genome-and transcriptome-level experiments. To get the correct runtimes for both tools, we excluded the execution time for a common step performed by an external tool (Bowtie, version 1) that prepares the index for the reference genome or transcriptome. For genome-level experiments, *PHASIS* displayed a minimum speed gain of 3x in Arabidopsis and rice and a maximum speed gain of 7x in maize **(Figure 3).** In transcriptome-level experiments, both tools took almost equal time **(Figure 3).** However, *PHASIS* yielded 10x (n=408) to 17x (n=9065) more *PHAS* predictions for 21-and 24*-PHAS* loci, respectively **(Table 2 and Supplementary Figure 3),** compared to *PhaseTank*, which means that *PHASIS* processed a high number of *PHAS* transcripts in the same runtime. Moreover, the time and effort required to convert the reference genome as well as the sRNA libraries to meet *PhaseTank* input requirements were not included in these runtime comparisons. Lastly, it should be noted that *PHASIS* takes significantly less time for any subsequent analyses in these species because of its unique ability to systematically store ancillary data in the first run, check data integrity and compatibility with parameters for subsequent runs, and avoid redoing the slowest steps, such as reference pre-processing, index preparation, etc.

**Figure 3.**
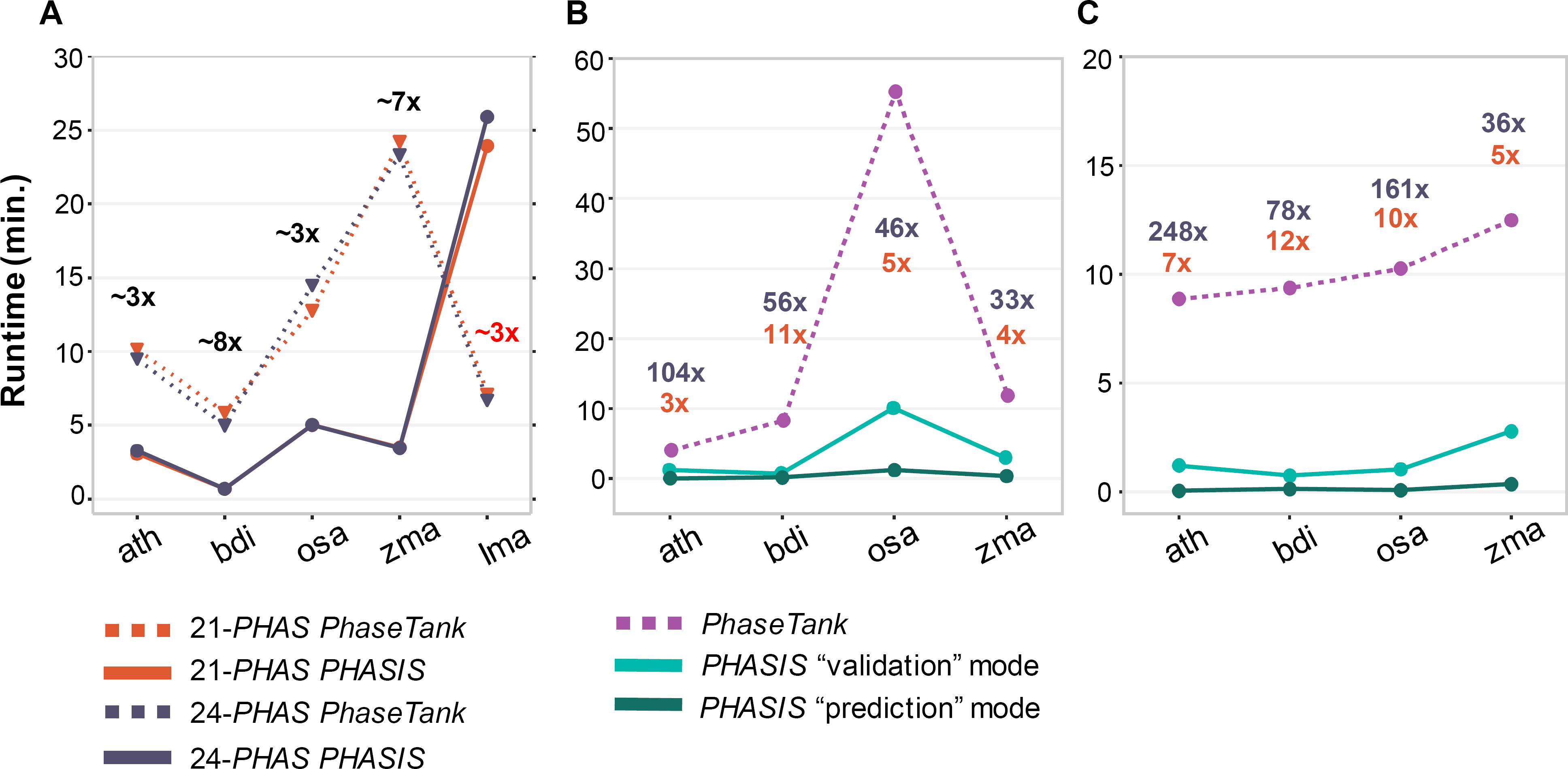
Runtime comparisons between *PHASIS* and *PhaseTank*. **A**) Time taken by both tools in prediction of **21-** and 24-*PHAS* loci or precursors transcripts. Speed gain displayed by *PHASIS* over *PhaseTank*, approximated for both size classes, is individually marked for each species. **B**) and **C**) Time taken by both tools in predicting **21-** and 24*-PHAS* triggers, respectively. Speed gain displayed by *PHASIS* in “validation” and “prediction” mode over *PhaseTank* is displayed in blue and orange colors respectively. In all comparisons, *Arabidopsis* is marked as “ath", Brachypodium as “bdi", rice as “osa", maize as “zma” and Lilium as “Ima".

### Comparison of PHASIS predictions with manually-curated data

We next wanted to address how well the predictions from *PHASIS* compare with a set of manually-curated *PHAS* loci. We and collaborators curated a set of 21-and 24*-PHAS* (n= 463 and 176 loci from precisely-staged, premeiotic and meiotic maize anthers [7]. This curated set was prepared by first combining all libraries from the sampled premeiotic and meiotic stages into a single file, followed by genome wide scans to identify phasiRNA generating loci using a score-based approach [5] and finally curating each *PHAS* locus to exclude those that overlap with repeat-associated regions or display sRNA distribution atypical of hc-siRNA generating loci [7]. *PHASIS* processes each library separately mainly to a) detect phased patterns independently in at least one of the input sRNA libraries, b) minimize any noise that could be added by combining sRNAs from multiple stages, tissues or treatments, and c) infer the correct 5’-end of *PHAS* loci by collating data from different libraries. Therefore, unlike the original analysis, we did not combine the 32 libraries (see **Supplementary Table** 1) for predictions by *PHASIS*. Furthermore, to emulate ‘real world’ conditions in which *PHASIS* would be used by non-experts, we did not provide a confidence cutoff - i.e. *PHASIS* was run in the default mode. Of the manually-curated 463 21*-PHAS* and 176 24-*PHAS* loci, *PHASIS* captured 89.0% (n=411) and 85.79% (n=151) **(Supplementary Table 5).** The majority of those missed either lacked continuous phased positions or had a very low abundance across all sRNA libraries, and some had a single sRNA read accounting for the major proportion (>90%) of the abundance at the *PHAS* locus. The average abundance of siRNAs in the ‘missed’ 21-and 24-*PHAS* set was ~12-and 252-times lower compared to the common pool (p < 1.02e-09), supporting the observation that those missed by *PHASIS* were weakly phased loci; a portion of these could be captured with a relaxed cutoff. Nonetheless, these results demonstrate that *PHASIS* predictions are largely consistent with the manually-curated data, and for most studies, the use of *PHASIS* may ameliorate the need to manually curate *PHAS* locus predictions, an otherwise complex and cumbersome task especially when *PHAS* loci number in the hundreds to thousands, as reported in many plant genomes [2,7,8,13,14,17].

### Trigger prediction and runtime performance

The identification of the miRNA triggers of *PHAS* loci is important for understanding their potential roles, classification and for discovery of secondary siRNA cascades. In addition, a set of *PHAS* loci or transcripts when combined with the trigger identity may serve as a gold-standard reference set for downstream experimental and bioinformatics studies. Given the importance of trigger identification, we compared the trigger prediction performance of *PHASIS* in ‘validation’ mode with *PhaseTank*. The *PHASIS* ‘validation’ mode will identify triggers for *PHAS* loci or transcripts using experimental data such as PARE, degradome or GMUCT libraries (‘PARE’, henceforth) [18-20]. *PhaseTank* by default predicts triggers in ‘validation’ mode, i.e. experimental data is required. Since *PHASIS* predicted more *PHAS* loci compared to *PhaseTank*, the number of *PHAS* loci (and transcripts) with the predicted triggers by *PHASIS* was higher too. So, for a fair comparison, we used only the common pool of *PHAS* loci to evaluate the trigger prediction performances. *PHASIS* displayed a gain of up to 76.0% in predicted triggers, except for 21*-PHAS* loci in Arabidopsis **(Figure 2A and B),** with a minimum accuracy of 96.0% for 24-*PHAS* maize loci and maximum accuracy of 99.5% in Brachypodium 21*-PHAS* loci **(Supplementary Table 3).** This accuracy was computed as the proportion of triggers (out of the total) that match to known triggers of phasiRNAs and tasiRNAs described in earlier studies [2,5,6,8,25-27]. These estimates of accuracy are likely conservative, given that there might be a few new and unknown triggers that we counted as false positives in our accuracy computations. We excluded rice 24*-PHAS* loci from our comparisons because both tools failed to report triggers for these loci, likely due to sRNA libraries that were not precisely staged relative to the accumulation of 24-nt phasiRNAs and thereby making it difficult to capture the 5’ and 3’ ends of *PHAS* loci – information crucial to the identification of correct triggers. Lilium 21- and 24-*PHAS* transcripts were also excluded from the comparisons because of a lack of PARE data from the corresponding anther stages, data required by *PhaseTank* to predict triggers. Likewise, Arabidopsis 24-*PHAS* couldn’t be included in our comparison as *PhaseTank* predicted loci (n=146) were false positives, and there were no overlapping loci with *PHASIS*.

We noticed a decline in number of predicted triggers by *PHASIS* for 21*-PHAS* loci in Arabidopsis, compared to those predicted by *PhaseTank* (Figure 2A). This decline in predicted triggers was traced to seven phased loci corresponding to the pentatricopeptide repeat (PPR) gene family, with phasiRNAs triggered by miR161. We found trigger sites predicted by *PhaseTank* for five of these loci, located 214 nt to 310 nt from the first or last phased cycle of the *PHAS* loci, towards their middle **(Supplementary Table 4).** Since *phastrigs*, the trigger discovery tool of *PHASIS*, is built with the aim to eliminate the need for experimental data and because trigger sites are expected to overlap with 5’ or 3’ ends of the phased region, it uses a narrow search space at the 5’ and 3’ ends to search for triggers. Hence, these miR161 target sites were missed by *PHASIS*. In *phastrigs*, the search space to identify triggers is defined by the number of phased positions (*PHAS*-index) on either side of 5’ and 3’ ends of phased regions, and by default *PHAS-*index is set to ± 3 positions for both ends. The *PHAS-index* setting to expand or the narrow search space for triggers is user tunable and can be adjusted to capture such cases. Nonetheless, these 21*-PHAS* loci from Arabidopsis support our estimates that trigger identification by *phastrigs* is conservative, and relaxing the *phastrigs* search parameters could further increase the gain in predicted triggers compared to *PhaseTank*.

### Identifying PHAS triggers without additional experimental data

We next evaluated the performance of *PHASIS* in trigger ‘prediction’ mode by comparing it with *PhaseTank* and *PHASIS* in the ‘validation’ mode. We define *PHASIS* ‘prediction’ mode as an analysis to predict triggers for *PHAS* loci or transcripts without any supporting experimental data such as PARE, degradome or GMUCT libraries. Lilium was excluded from the comparison of predicted triggers due to the lack of PARE data, which is compulsory for *PhaseTank* to predict triggers and required by *PHASIS* in ‘validation’ mode. Also, for reasons mentioned above, 24-*PHAS* loci from Arabidopsis and rice were excluded from the comparisons. *PHASIS* displayed a minimum gain of 40.3% and maximum gain of 178.3% over *PhaseTank* in predicting triggers for 21*-PHAS* and 24*-PHAS* loci from Brachypodium, respectively **(Table 2** and **Figure 2).** The gain in the number of triggers ranged from a minimum of 35 for maize 24-*PHAS* loci to a maximum of 611 for rice 21*-PHAS* loci. In addition to the gain in trigger prediction, *PHASIS* also displayed significant accuracy in prediction mode, with a minimum accuracy of 89.9% in predicting triggers for 24-*PHAS* loci from maize and maximum accuracy of 99.9% in the case of Lilium 24-*PHAS* precursor transcripts, however, with an exception for Lilium 21*-PHAS* triggers. The accuracy of predicted triggers of Lilium 21*-PHAS* loci was significantly lower (43.9%) compared to the other species **(Supplementary Table 2).** For Lilium, we used miRNAs from well-characterized monocots like rice and maize because a complete set of miRNAs were not available due to the absence of a sequenced genome. Surprisingly, we found that for Lilium 21-*PHAS* transcripts a majority of triggers corresponded to miR2275 instead of miR2118; this observation was puzzling because miR2275 is known to trigger 24-nt phasiRNAs in the grasses [2,7,13], and this was the basis for the low recorded accuracy in predicting Lilium 21*-PHAS* triggers. We did not further investigate the miR2275-triggered 21*-PHAS* transcripts. We also noticed that the proportion of 21-and 24-*PHAS* precursors for which triggers could be identified in Lilium, 18.1% and 25.9% respectively **(Table 2 and Figure 2),** was substantially lower compared to the overall average of 73.8% in other species for which genomic analysis was performed. Plant *PHAS* precursor transcripts are typically cleaved by the miRNA trigger, converted to dsRNA by an RNA-dependent RNA polymerase, and then successively diced by a Dicer enzyme. Since no data on transcriptional rate, stability and half-life of phasiRNA precursors are available, we speculated that a portion of the Lilium *PHAS* precursor transcripts were shortened by processing from the 5’ end, removing the trigger target sites. Identifying triggers from such “processed” precursor transcripts is not possible because the P1 site corresponding to the first phasiRNA (at the 5’ terminus) could be missing from the transcript. In addition, the presence of already-processed mRNAs will confound the *de novo* assembly of precursor transcripts from short-reads.

To test whether the low yield of triggers by *phastrigs* resulted from our use of processed precursor transcripts and not a technical shortcoming of *PHASIS*, we generated Single Molecule Real Time (SMRT) PacBio sequencing data from Lilium anthers 4 mm to 6 mm in length. These sizes represented premeiotic and meiotic stages of anther development **(see supplementary methods)** and were selected based on the availability of the samples. Capturing *PHAS* precursors is complex, not just because these are targets of miRNAs presumably rapidly processed by a Dicer, but reproductive phasiRNAs are ephemeral in development and thus not easily captured [7]. SMRT-seq produced 425,897 full-length transcripts for 176,373 unique isoforms, which were pre-processed to generate 122,779 high quality (polished) transcripts. This set had 5,131 unique proteins covered by more than 80% protein length, relative to the Uniprot protein-sequence resource, thereby suggesting a reasonable assembly of the anther transcriptome. *PHASIS* identified 87 21*-PHAS* and 175 24-*PHAS* precursor transcripts. This low yield of *PHAS* transcripts was expected, though not to such a degree, because of the combination of the following: a) low read counts for SMRT-seq compared to the deep RNA-seq data, b) the coverage-based error correction algorithm - ‘Quiver’ implemented in the IsoSeq protocol (SMRT Analysis software version 2.3, Pacific Biosciences) which filters out transcripts with insufficient coverage, i.e. those that cannot be confidently corrected, and c) the aforementioned processive cleavage of *PHAS* precursors by Dicer. *phastrigs* could identify triggers for only 21.8% (n=19) of 21*-PHAS* precursors, a slight increase compared to 18.1% in the RNA-seq assembly, and these triggers included miR2275, miR2118 and miR390. This low proportion of triggers detected for 21*-PHAS* could result from missing the precise stage at which 21*-PHAS* precursors accumulate in the Lilium samples. However, *phastrigs* could identify triggers for 54.2% of the 24-*PHAS* precursors, a significant increase over the 25.9% in the RNA-seq assembly, supporting our premise about the completeness of the *PHAS* precursor transcripts. The processed precursors were likely collapsed into the full-length or the longest transcript in SMRT-seq assembly, thereby enriching the proportion of uncleaved precursor transcripts. Hence, it should be noted that neither the precursors from neither RNA-seq nor SMRT-seq may accurately represent the true total count of *PHAS* loci in *Lilium*.

Lastly, we compared runtimes for both tools for miRNA trigger prediction of *PHAS* loci and transcripts. *PHASIS* showed a minimum speed gain of 3.3x and a maximum speed gain of 12.6x over *PhaseTank* in ‘validation’ mode **(Figure 3).** In ‘prediction’ mode, *PHASIS* was at least 5.0x and at most 31.2x faster compared to its own ‘validation’ mode without any significant loss in accuracy (Supplementary Table 3). *PhaseTank* requires PARE data to predict triggers, and lacks a function equivalent to *PHASIS* ‘prediction’ mode, but since *PHASIS*, even without the additional experimental data (like PARE) displays >89.9% accuracy in trigger prediction, we decided to compare runtimes for both. *PHASIS* in ‘prediction’ mode displayed a minimum speed gain of 33.3x and a maximum gain of 104.3x for Arabidopsis 21*-PHAS* loci **(Figure 3).** The trigger predictions for 24-*PHAS* loci from Arabidopsis and rice, which displayed even higher speed gains, were excluded from the runtime comparisons due to the reasons described above. This gain in *PHAS* trigger identification demonstrates the capacity of *PHASIS* to predict triggers without experimental data. This functionality will save time and the cost of preparing PARE libraries; it will also reduce the amount of sample required for phasiRNA analysis. Protocols for preparing PARE libraries require comparatively more input RNA relative to RNA-seq or sRNA-seq [28].

## Conclusions

Loci generating 21-and 24-nt phasiRNAs are widely prevalent across land plants [2,5,8,14,16,17,29], varying in numbers per genome from tens to thousands, displaying diverse spatial and temporal expression patterns, and participating in an array of different functions [5,7,8,29,30]. Recently, piRNAs in *Drosophila* too were reported to be phased, generating ‘trailer’ piRNAs in 27-nt intervals after cleavage by secondary siRNA and Zucchini-dependent processing of cleaved transcript [21,22]. Given the wide prevalence of phasiRNAs and the rate of genome sequencing, it is likely that they will be better characterized and studied in the coming years. The existing tools for computational characterization of *PHAS* loci or transcript are limited both in number and functionality.

The *PHASIS* suite provides an integrated solution for the large-scale survey of tens to hundreds of sRNA libraries for the following applications: a) *de novo* discovery of *PHAS* loci and precursor transcripts, b) a summarization of *PHAS* loci from specific groups of sRNA libraries, c) a comparison of *PHAS* summaries between groups corresponding to samples from different stages, tissues and treatments, d) quantification and annotations of *PHAS* loci, and e) discovery of their miRNA triggers. *PHASIS* generates easily parsed output files for downstream bioinformatics analysis, formatted result files for immediate consumption and organized ancillary data to facilitate optimizations like a re-summarization to exclude or include libraries.

More complete characterization of phasiRNAs in evolutionarily diverse plant genomes will advance our understanding of phasiRNA function and the adaptation of the pathway, and it may yet discover new classes of *PHAS* genes. *PHASIS* will thus facilitate the discovery of phasiRNAs and their precursors, and the identification of their triggers by eliminating the requirement of a genome assembly and experimental PARE/degradome data. *PHASIS* offers flexibility to users to tailor analyses for their own goals and it integrates an array for functions in one package.

## Methods

*PHASIS* comprises three components that together perform *de novo* discovery, annotation, quantification, comparison and trigger identification for *PHAS* loci or precursor transcripts. We chose a modular approach over the single ‘one-commanď style for the following reasons: a) to maximize the flexibility for specific data or study requirements; b) to integrate multiple, connected analyses; and, c) to reduce overall runtime by maximizing phase- and step-specific parallelization. A description of these tools-*phasdetect, phasmerge, phastrigs* – in order of their utility or phases of study is provided below (see also **Figure 1**). *PHASIS* leverages the Python (v3) process-based “threading” interface to achieve efficient scalability and significantly reduce runtimes through parallel computing.

*phasdetect* performs *de novo* prediction of *PHAS* loci or precursor transcripts using user-supplied sRNA libraries along with a reference genome or transcriptome. It can efficiently process tens to hundreds of sRNA libraries in parallel, reducing runtimes. *phasdetect* operates via three main steps: a) first, sRNA libraries are normalized and mapped to the reference; b) second, mapped sRNA reads are scanned to identify regions rich for specific size classes, such as those generated by Dicer activity (typically 21, 22, or 24 nt in plants); and, c) finally these regions are stitched into clusters and the phasing of the small RNAs is computed as a *p-value*. We adopted a standard approach to compute *p-values* [9]. Parameters controlling these steps can be modified by users via the setting file “phasis.set”, including values for *phase, mindepth* and *clustbuffer;* these refer to the phasing periodicity, minimum sRNA abundance to be included for *p-value* computation, and the minimum distance separating two clusters. These parameters are explained in detail on the *PHASIS* wiki page (https://github.com/atulkakrana/PHASIS/wiki/). The output for *phasdetect* includes library-specific list of *PHAS* loci (or transcripts) at several different confidence levels plus ancillary data, used to reduce runtime for subsequent analyses. For example, in case of a reanalysis after adding new libraries, *phasdetect* checks for any changes in parameters from the earlier analysis, assesses the integrity and compatibility of the ancillary data for, and reuses existing data to avoid repetition. This ancillary data also enables an array for downstream analyses and analysis-specific optimizations directly through *phasdetect*.

*phasmerge* generates a summary, matches *PHAS* loci to annotations and performs a comparison between the *PHAS* summaries using the library-specific *PHAS* lists and ancillary data generated by *phasdetect*. These operations are selected by using the *-mode* option with the *merge* (default) or *compare* values. The *merge* mode prepares a *PHAS* summary for the libraries of interest, or for libraries that belong to different groups based on sample stages, tissues or treatments. The analysis can be tailored to meet the study requirements. For example, to maximize discovery, a user might set a lower confidence level (*p-value*) for summarization and consider all loci with a trigger predicted without the PARE data (identified through *phastrigs)* for downstream analyses. In contrast, a user motivated to maximize the quality might identify *PHAS* loci with the highest confidence level, followed by pruning of results with stringent quality parameters (described on the *phasmerge* wiki), and use *PHAS* loci that have PARE-supported triggers. *PHAS* summaries from different groups of libraries can be compared using *compare* mode. This is particularly useful to identify intersecting and exclusive *PHAS* loci between different groups of stages, tissues or treatments. In *merge* mode, if an additional annotation file is provided, then merged *PHAS* loci are matched to genome annotations so as to identify coding *PHAS* loci or other available annotations. This function also supports quick discovery of precursor transcripts for summarized *PHAS* loci when provided with a GTF file generated from mapping the transcriptome assembly to genome. Furthermore, *phasmerge* attempts to determine the correct 5’ terminus of *PHAS* loci by optimizing for the best 5’ or 3’ coordinates based on the user’s sRNA data – a crucial functionality for determination of the correct miRNA trigger. *phasmerge* benefits from the modular *PHASIS* workflow, allowing users to optimize their results for the study which may vary in purpose, and making *phasmerge* independent from other tools.

The *phasmerge* workflow has three mandatory and two optional steps: a) via *merge* mode, *phasmerge* first generates a unique list of *PHAS* loci (or transcripts) for each user-specified library, by selecting predictions with the highest available confidence score (lowest *p-value)* that pass a user-supplied *p-value* cutoff, after comparing predictions from all available confidence levels; b) *phasmerge* clusters the “best” candidate loci from specified libraries specific by the user, based on the degree of overlap in phased positions (or ‘cycles’) to select a representative locus for each cluster; finally, c) *phasmerge* computes library-specific abundances, a size-class ratio, the maximum to total phasiRNAs abundance ratio, and other quality information. Optional steps include d) *compare* mode, which first reads *PHAS* loci (or transcripts) from user-supplied summaries (n=2) and then identifies matching *PHAS* pairs based on the overlap in phased positions, to report a combined matrix including both shared and unique loci in each *PHAS* summary file, and e) *merge* mode; when supplied with annotations, as described above, *phasmerge* matches a merged set of *PHAS* loci with genome annotations or with a genome-matched transcriptome assembly, both provided as GTF file, to report exonic or complete overlaps with annotated transcripts. This step requires prior installation of *SQLite* on user’s machine. *phasmerge* generates several reports as output, most importantly, *PHAS* summary for libraries of interest which includes quality parameters (see online wiki for more information), FASTA files for size-specific siRNAs and all the siRNAs from phased positions along with detailed information on phased clusters with phasiRNAs, positions, associated *p-values*, etc.

*phastrigs* identifies sRNA triggers for *PHAS* loci and precursor transcripts using the *phasmerge* summaries and a user-provided list of miRNAs (or any other small RNA). It was developed with the idea to minimize the requirement of experimental PARE libraries [18-20]. However, if such data (‘PARE’, henceforth) are provided, then *phastrigs* reports sRNA triggers with experimental support; these may be of higher confidence for some downstream experimental analyses.

*phastrigs* uses an algorithm designed to be both fast and exhaustive. It uses *miRferno*, an exhaustive target prediction algorithm that we developed [31] to predict target sites for user-supplied miRNAs. The speed and precision of *phastrigs* is enhanced by a scan focused on the 5’ terminus of each *PHAS* locus (5’-end of the first cycle, the P1 position) for the trigger site, which reduces the search space and chance of reporting false triggers. This 5’ terminus is inferred at the summarization step by *phasmerge* while collating data from different sRNA libraries. In the case of *PHAS* transcripts, only the 5’ terminus of the phased precursor is scanned, while in case of genomic *PHAS* loci, either the 5’ or 3’ end of the phased region is chosen, based on the strand targeted by a specific miRNA. *phastrigs* analysis is divided into two main steps: a) *PHAS* transcripts or genomic sequences are extracted, and targets for user-supplied miRNAs are predicted; b) next, a scan of phased positions located at the 5’ or 3’ termini of precursor for a target site that corresponds with the production of phasiRNAs is performed; this scan looks for target sites within ± 3 nt of the *‘PHAS* index’, defined as theoretical phased positions upstream from the 5’ terminus of P1. If PARE data is supplied, then PARE-validated cleavage sites are used for trigger identification. The *phastrigs* report includes detailed information on miRNA-target interactions, PARE abundances at the predicted cleavage site, and the *PHAS* index of the predicted trigger site relative to the P1 position.

### Software

The methods and algorithm described in this article, implemented as *PHASIS* suite of tools for *PHAS* discovery, are freely available from https://github.com/atulkakrana/PHASIS. *PHASIS* is released under the OSI Artistic License 2.0. Tools and Perl libraries required to use *PHASIS* along with the instructions to install and usage of individual tools is provided in detail in the *PHASIS* wiki (https://github.com/atulkakrana/PHASIS/wiki/).

## Additional files

**Additional file 1:** Supplementary methods in Microsoft Word (.doc) format.

**Additional file 2:** Supplementary figures from S1 to S3 in Portable Document Format (PDF).

**Additional file 3:** Supplementary tables from S1 to S4 in Office Open XML Spreadsheet format (.xlsx)

## Abbreviations

*PHASIS*: *PHAS* Inspection Suite
tasiRNAs: *trans-*acting siRNAs
PMC: pollen mother cells
PARE: Parallel Analysis of RNA Ends
SMRT: Single Molecule Real Time Sequencing
GMUCT: genome-wide mapping of uncapped and cleaved transcripts

## Acknowledgments

We thank Bruce Kingham and Olga Shevchenko from Delaware Biotechnology Institute (Newark, DE, USA) for their help with PacBio sequencing. We thank Karol Miaskiewicz from Delaware Biotechnology Institute (Newark, DE, USA) for work on the SMRT portal and the *PB-Tofu* command line environment. We also thank Kun Huang from University of Delaware (Newark, DE, USA) for helpful discussions on the developmental stage and anther size correlations.

### Funding

This work was supported by U.S. National Science Foundation Plant Genome Research Program (NSF-PGRP) grant (1649424) and University Competitive Fellow Award (2015-2016) from University of Delaware.

### Author contributions

AK conceived the project. AK and PL designed and implemented the method, individual contributions are marked on scripts. PP, RH, DA and AK tested tools and compiled benchmarking data. AK and SM collected the Lilium samples. SM prepared the SMRT sequencing libraries. AK and BCM wrote the manuscript. All authors read and approved the final manuscript.

### Competing interests

The authors declare that they have no competing interests.

## References

1. Axtell MJ. Classification and comparison of small RNAs from plants. Annu. Rev. Plant Biol. 2013;64:137–59.

2. Johnson C, Kasprzewska A, Tennessen K, Fernandes J, Nan G-L, Walbot V, et al. Clusters and superclusters of phased small RNAs in the developing inflorescence of rice. Genome Res. 2009;19:1429–40.

3. Vazquez F, Vaucheret H, Rajagopalan R, Lepers C, Gasciolli V, Mallory AC, et al. Endogenous trans-Acting siRNAs Regulate the Accumulation of Arabidopsis mRNAs. Mol. Cell. 2004;16:69–79.

4. Peragine A, Yoshikawa M, Wu G, Albrecht HL, Poethig RS. SGS3 and SGS2/SDE1/RDR6 are required for juvenile development and the production of trans-acting siRNAs in Arabidopsis. Genes Dev. 2004;18:2368–79.

5. Allen E, Xie Z, Gustafson AM, Carrington JC. microRNA-directed phasing during trans-acting siRNA biogenesis in plants. Cell. 2005;121:207–221.

6. Fei Q, Xia R, Meyers BC. Phased, secondary, small interfering RNAs in posttranscriptional regulatory networks. Plant Cell. 2013;25:2400–15.

7. Zhai J, Zhang H, Arikit S, Huang K, Nan G-L, Walbot V, et al. Spatiotemporally dynamic, cell-type-dependent premeiotic and meiotic phasiRNAs in maize anthers. Proc. Natl. Acad. Sci. U. S. A. 2015;112:3146–51.

8. Zhai J, Jeong D-H, Paoli ED, Park S, Rosen BD, Li Y, et al. MicroRNAs as master regulators of the plant NB-LRR defense gene family via the production of phased, trans-acting siRNAs. Genes Dev. 2011;25:2540–53.

9. Chen H-M, Chen L-T, Patel K, Li Y-H, Baulcombe DC, Wu S-H. 22-Nucleotide RNAs trigger secondary siRNA biogenesis in plants. Proc. Natl. Acad. Sci. U. S. A. 2010;107:15269–74.

10. Cuperus JT, Carbonell A, Fahlgren N, Garcia-Ruiz H, Burke RT, Takeda A, et al. Unique functionality of 22-nt miRNAs in triggering RDR6-dependent siRNA biogenesis from target transcripts in Arabidopsis. Nat. Struct. Mol. Biol. 2010;17:997–1003.

11. Jouannet V, Moreno AB, Elmayan T, Vaucheret H, Crespi MD, Maizel A. Cytoplasmic Arabidopsis AG07 accumulates in membrane-associated siRNA bodies and is required for ta-siRNA biogenesis. EMBO J. 2012;31:1704–13.

12. Li S, Le B, Ma X, Li S, You C, Yu Y, et al. Biogenesis of phased siRNAs on membrane-bound polysomes in Arabidopsis. eLife. 2016;5:e22750.

13. Jeong DH, Schmidt SA, Rymarquis LA, Park S, Ganssmann M, German MA, et al. Parallel analysis of RNA ends enhances global investigation of microRNAs and target RNAs of Brachypodium distachyon. Genome Biol. 2013;14:R145.

14. Arikit S, Xia R, Kakrana A, Huang K, Zhai J, Yan Z, et al. An atlas of soybean small RNAs Identifies phased siRNAs from hundreds of coding genes. Plant Cell. 2014;26:4584–601.

15. Xia R, Ye S, Liu Z, Meyers BC, Liu Z. Novel and recently evolved microRNA clusters regulate expansive F-BOX gene networks through phased small interfering RNAs in wild diploid strawberry. Plant Physiol. 2015;169:594–610.

16. Fei Q, Yang L, Liang W, Zhang D, Meyers BC. Dynamic changes of small RNAs in rice spikelet development reveal specialized reproductive phasiRNA pathways. J. Exp. Bot. 2016;67:6037–49.

17. Xia R, Xu J, Arikit S, Meyers BC. Extensive families of miRNAs and *PHAS* Loci in Norway Spruce demonstrate the origins of complex phasiRNA networks in seed plants. Mol. Biol. Evol. 2015; 32: 2905–2918.

18. German MA, Luo S, Schroth G, Meyers BC, Green PJ. Construction of Parallel Analysis of RNA Ends (PARE) libraries for the study of cleaved miRNA targets and the RNA degradome. Nat. Protoc. 2009;4:356–62.

19. Addo-Quaye C, Eshoo TW, Bartel DP, Axtell MJ. Endogenous siRNA and miRNA targets identified by sequencing of the Arabidopsis degradome. Curr. Biol. 2008;18:758–762.

20. Gregory BD, O’Malley RC, Lister R, Urich MA, Tonti-Filippini J, Chen H, et al. A link between RNA metabolism and silencing affecting Arabidopsis development. Dev. Cell. 2008;14:854–866.

21. Mohn F, Handler D, Brennecke J. piRNA-guided slicing specifies transcripts for Zucchini-dependent, phased piRNA biogenesis. Science. 2015;348:812–7.

22. Han BW, Wang W, Li C, Weng Z, Zamore PD. piRNA-guided transposon cleavage initiates Zucchini-dependent, phased piRNA production. Science. 2015;348:817–21.

23. Guo Q, Qu X, Jin W. PhaseTank: genome-wide computational identification of phasiRNAs and their regulatory cascades. Bioinformatics. 2015;31:284–6.

24. Axtell MJ. ShortStack: Comprehensive annotation and quantification of small RNA genes. RNA. 2013;19:740–51.

25. Allen E, Howell MD. miRNAs in the biogenesis of trans-acting siRNAs in higher plants. Semin. Cell Dev. Biol. 2010;21:798–804.

26. Axtell MJ, Jan C, Rajagopalan R, Bartel DP. A two-hit trigger for siRNA biogenesis in plants. Cell. 2006;127:565–77.

27. Zheng Y, Li Y-F, Sunkar R, Zhang W. SeqTar: an effective method for identifying microRNA guided cleavage sites from degradome of polyadenylated transcripts in plants. Nucleic Acids Res. 2012;40:e28.

28. Zhai J, Arikit S, Simon SA, Kingham BF, Meyers BC. Rapid construction of parallel analysis of RNA end (PARE) libraries for illumina sequencing. Methods. 2014;67:84–90.

29. Shivaprasad PV, Chen H-M, Patel K, Bond DM, Santos BACM, Baulcombe DC. A microRNA superfamily regulates nucleotide binding site-leucine-rich repeats and other mRNAs. Plant Cell. 2012;24:859–74.

30. Dukowic-Schulze S, Sundararajan A, Ramaraj T, Kianian S, Pawlowski WP, Mudge J, et al. Novel meiotic miRNAs and indications for a role of phasiRNAs in meiosis. Plant Genet. Genomics. 2016;762.

31. Kakrana A, Hammond R, Patel P, Nakano M, Meyers BC. sPARTA: a parallelized pipeline for integrated analysis of plant miRNA and cleaved mRNA data sets, including new miRNA target-identification software. Nucleic Acids Res. 2014;42:el39.

